# Chloride binding to prestin does not influence very high-frequency complex nonlinear capacitance (cNLC) in the mouse outer hair cell

**DOI:** 10.1101/2024.01.29.577264

**Authors:** Jun-Ping Bai, Chenou Zhang, Iman Bahader, Nicola Strenzke, Vijay Renigunta, Dominik Oliver, Dhasakumar Navaratnam, Oliver Beckstein, Joseph Santos-Sacchi

## Abstract

Prestin (SLC26a5) function evolved to enhance auditory sensitivity and frequency selectivity by providing mechanical feedback via outer hair cells (OHC) into the organ of Corti. Its effectiveness is governed by the voltage-dependent kinetics of the protein’s charge movements, namely, nonlinear capacitance (NLC). We study the frequency response of NLC in the mouse OHC, a species with ultrasonic hearing. We find that the characteristic frequency cut-off (F_is_) for the mouse in near 27 kHz. Single point mutations within the chloride binding pocket of prestin (e.g., S396E, S398E) lack the protein’s usual anion susceptibility. In agreement, we now show absence of anion binding in these mutants through molecular dynamics (MD) simulations. NLC F_is_ in the S396E knock-in mouse is unaltered, indicating that high frequency activity is not governed by chloride, but more likely by viscoelastic loads within the membrane. We also show that the allosteric action of chloride does not underlie piezoelectric-like behavior in prestin, since tension sensitivity of S396E NLC is comparable to that of WT. Because prestin structures of all species studied to-date are essentially indistinguishable, with analogous chloride binding pockets, auditory requirements of individual species for cochlear amplification likely evolved to enhance prestin performance by modifying, not its protein-anion interaction, but instead external mechanical loads on the protein.

**Significance:** Prestin is believed to provide cochlear amplification in mammals that possess a wide range of frequency sensitivities. Previously, chloride anions have been shown to control prestin kinetics at frequencies below 10 kHz. However, now we find that chloride binding is not influential for prestin kinetics in the very high range of the mouse. We suggest that such high frequency prestin performance is governed by impinging mechanical loads within the membrane, and not interactions with anions.

## Introduction

The mammalian organ of Corti houses two types of sensory receptor cells, the inner (IHC) and outer hair cells (OHC). Each is responsive to sound stimuli, producing receptor potentials that are tuned to particular acoustic frequencies, depending upon the location of those cells along the tonotopic cochlear spiral (Russell and Sellick, 1978; Dallos et al., 1982). Unlike the IHC, the OHC additionally is mechanically active, producing changes in length (electromotility, **eM**) when electrically stimulated (Brownell et al., 1985). OHC **eM**, being voltage dependent (Santos-Sacchi and Dilger, 1988), is the basis of receptor potential-driven amplificatory feedback, termed cochlear amplification, that enhances auditory response sensitivity and selectivity of the IHC, the main sensory cell type. The voltage-dependent protein, prestin (Zheng et al., 2000), which resides exclusively in the OHC lateral membrane, underlies this process (Dallos et al., 2008). It is not surprising that chloride anions play a central role in OHC/prestin activity (Oliver et al., 2001; Santos-Sacchi et al., 2006a; Rybalchenko and Santos-Sacchi, 2008), since prestin (SLC26a5) is a member of the SLC26 anion transporter family (Zheng et al., 2000; Geertsma and Oliver, 2023).

The mechanical activity of prestin arises from its voltage-sensor charge movement, traditionally measured as a robust nonlinear, bell-shaped, voltage-dependent capacitance (NLC) (Santos-Sacchi, 1991), which drives conformational switching of the protein. This activity across frequency is limited by the protein’s kinetics within the plasma membrane (Santos-Sacchi and Tan, 2018). The characteristic speed of a voltage-dependent protein is identified at the membrane voltage (V_h_) where sensor charge (Q) is equally distributed across the membrane field, namely, at the midpoint of its Q-V Boltzmann function or, correspondingly, at the peak of its NLC. Interestingly, prestin is also sensitive to membrane tension, shifting V_h_ and generating displacement currents upon membrane stretch, observations that have led to its characterization as piezoelectric-like (Iwasa, 1993; Gale and Ashmore, 1994; Kakehata and Santos-Sacchi, 1995; Takahashi and Santos-Sacchi, 2001; Santos-Sacchi and Tan, 2022). Though the frequency response of peak NLC under voltage clamp in guinea pig OHC membrane patches has a characteristic frequency roll-off near 10-20 kHz (Gale and Ashmore, 1997; Dong et al., 2000; Santos-Sacchi and Tan, 2019; Santos-Sacchi and Tan, 2022), significant activity is measurable in the ultrasonic range (Santos-Sacchi et al., 2023)..

Apart from piezoelectric-like influences on prestin frequency response (Santos-Sacchi and Tan, 2022), we also find that chloride anions can influence prestin kinetics (Santos-Sacchi and Song, 2016; Santos-Sacchi and Tan, 2022), with mutations that affect anion binding site access having measurable effects (Takahashi et al., 2016; Takahashi et al., 2023). These effects have been demonstrated in the low frequency range (<10kHz). Since it is unlikely that prestin tertiary structure differs significantly among species, with gerbil, human and dolphin already determined (Bavi et al., 2021; Ge et al., 2021; Butan et al., 2022; Futamata et al., 2022), we have hypothesized that chloride concentration could vary along the cochlea spiral or among mammals to influence kinetics to match species-specific acoustic requirements (Santos-Sacchi and Tan, 2022). Here we extend our physiological studies to the mouse, a species with acoustic frequency sensation above that of the guinea pig, to investigate the influence of chloride binding on the high frequency kinetics of prestin. We utilize a prestin knock-in (KI) mouse (S396E), which lacks prestin anion sensitivity. Additionally, we complement our investigations with molecular dynamics simulations. We find that the ability to bind chloride does not influence high frequency NLC. Given the stretched, multi exponential nature of prestin frequency response, our data suggest that viscoelastic characteristics of the plasma membrane predominantly limit prestin’s high frequency kinetics.

## Materials and Methods

### Nonlinear Capacitance

Control OHC complex NLC (cNLC) was measured in b6 mouse (Jackson Lab; 1-2 months old) macro-patches under voltage clamp, with current sampling at 1 MHz (16 bit NI-USB 6356; National Instruments) using an Axon 200B amplifier in capacitive feedback mode and Bessel filter set to 100 kHz. Analyses were made in jClamp (www.scisoftco.com) and Matlab (www.mathworks.com). Experimental methods for NLC assessment are fully detailed in (Santos-Sacchi et al., 2021; Santos-Sacchi et al., 2023). Briefly, extracellular solution was (in mM): NaCl 100, TEA-Cl 20, CsCl 20, CoCl_2_ 2, MgCl_2_ 1, CaCl_2_ 1, Hepes 10, pH 7.2, and this solution was used in the on-cell patch pipette. Macro-patches were established on the OHC lateral plasma membrane, and swept-frequency AC voltage chirps (10 mV) were presented atop DC holding potentials ranging from +160 to -160 mV (see **Fig. 1** in (Santos-Sacchi et al., 2023)). The voltage chirps were generated in jClamp using the Matlab logarithmic “chirp” function (10 mV pk; pts = 4096; F0 = 244.141 Hz; F1 = 500 kHz; t1 = 0.004095 s). Currents were corrected for system roll-off characteristics in the frequency domain. The methodology of Fernandez et al. (Fernandez et al., 1982) was used to extract real and imaginary components of cNLC from membrane admittance, *Y*_*m*_(ω), as we previously described in detail (Santos-Sacchi et al., 2023). The complex components represent voltage sensor charge moving in phase (Im(cNLC)) and 90° out of phase (Re(cNLC)) with AC voltage, the former membrane-constrained charge movements being “resistive” and the latter being “capacitive” in nature. cNLC estimates were obtained following subtraction of linear admittance at +160 mV holding potential, where OHC NLC is absent, thereby removing stray capacitance effects. Currents were averaged 64 times, and spectra were obtained by FFT using 4096 data points.

**Figure 1.**
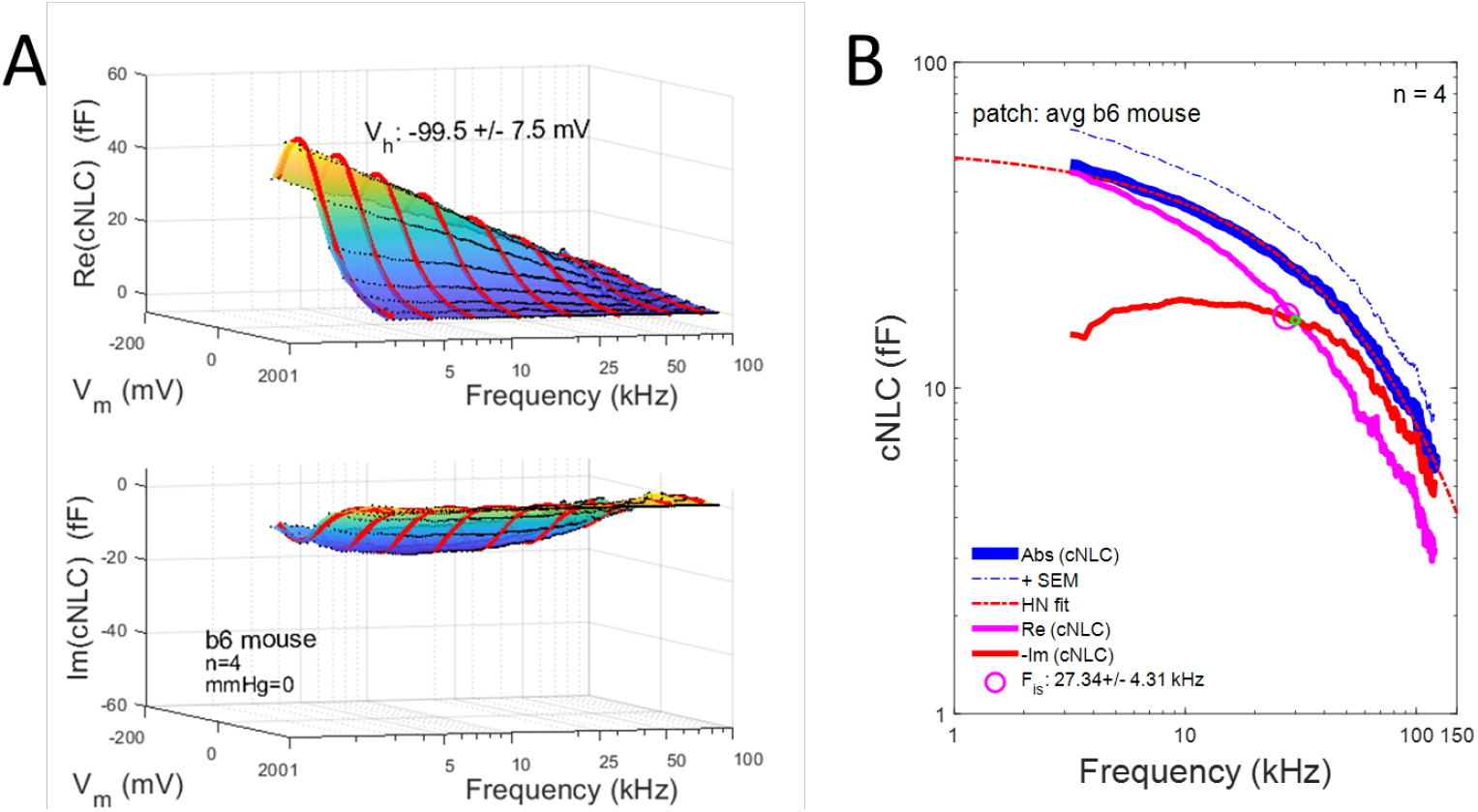
Average complex NLC in control b6 mouse OHC membrane patches. **A)** Real (top panel) and imaginary (bottom panel) components of cNLC across frequency and membrane voltage. Black lines indicate data and red lines are Boltzmann fits at selected frequencies. Averages (SE) @ 5.12 kHz V_h_ : -99.5 (7.5) mV; Q_max_: 7.6 (1.9) fC; *z*: 0.55 (0.03). Note that real component decreases with frequency while imaginary component increases then decreases with frequency. This pattern is like that of the guinea pig. No pipette pressure is delivered. **B)** Average frequency response of cNLC components near V_h_. Abs (cNLC) decreases across frequency in a stretched-exponential manner, indicated by the HN pink dotted line fit. The contribution of the imaginary component to absolute magnitude dominates at the highest frequencies. Average F_is_ (magenta circle) determined from individual recordings is 27.34 (4.31) kHz, faster than that measured in guinea pigs (Santos-Sacchi et al., 2023). Green point marker indicates intersection of averaged traces at 30.0 kHz.

NLC was fit with a Boltzmann function (Santos-Sacchi, 1991; Santos-Sacchi and Navarrete, 2002).

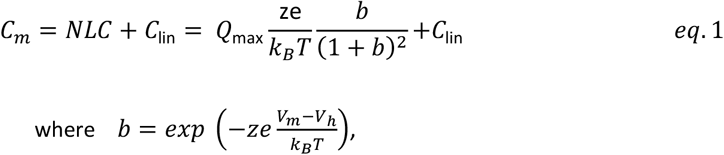

Q_max_ is the maximum nonlinear charge moved, V_h_ is voltage at peak capacitance or equivalently, at half-maximum charge transfer, V_m_ is membrane potential, *z* is valence, C_lin_ is linear membrane capacitance, e is electron charge, *k*_*B*_ is Boltzmann’s constant, and T is absolute temperature. Experiments were performed at room temperature. Fits were made to real and imaginary components of **cNLC** across interrogation frequency. Reported NLC averages were made at 5.12 kHz on Re(cNLC). No junction potentials are expected for the symmetrical pipette/extracellular solutions. Average zero current voltages of the membrane patches, which correspond to cell potentials (Mason et al., 2005), were -0.72, and +5.12 mV for the b6 control, and S396E prestin OHCs, respectively, were used to correct V_h_ values.

As previously (Santos-Sacchi et al., 2023), we fit the Abs(cNLC) data with the absolute value of the Havriliak-Negami [HN] relaxation function (Alverez et al., 1991), an empirical frequency domain representation of a stretched exponential function, only for the purpose of quantifying magnitude roll-off.

Cut-off frequencies (F_is_) were determined from the intersection of real and imaginary components of individual patches (magenta circles, see Results). Additionally, intersections of real and imaginary components for the plotted averaged NLC traces are denoted in the plots by a green marker. Means and standard errors of the mean (SEM) are reported in figures and legends. Two tailed T-tests were made in Excel. Methods have been approved by Yale’s animal use committee.

### S396E knock-in mouse

The S396E knock-in (KI) mouse was generated in D. Oliver’s lab, and details beyond those presented here are in preparation (Oliver et al., in prep). Briefly, the mPrestin-S396E mice were generated by CRISPR/Cas technology. Cas9 and guide RNA were delivered as RNPs along with a single-stranded DNA (ssDNA) repair template into mouse zygotes by pronuclear injection. The injected zygotes were cultured and allowed to develop to the 2-cell stage before being transferred into recipient C57Bl/6N females. Successful and selective introduction of the single point mutation was verified by sequencing the entire SLC26A5 gene. Mice were backcrossed for > 3 generations to C57Bl/6 mice. ABR and DPOAE were measured essentially as described in (Jing et al., 2013), except for a technical upgrade to the system to TDT System III (Tucker-Davis Technologies) with MF1 speakers (TDT) for DPOAE recordings. ABRs were measured at the age of 13 weeks. ABR and DPOAE recordings were approved by the local committee and the authorities of the state of Lower Saxonia, Germany under protocol number 33.19-42502-04-19/3123.

### Molecular Dynamics Simulations

#### Model preparation

We performed model simulations that were assembled based on our solved structure from gerbil prestin (Butan et al., 2022), namely, PDB: 7SUN and the Alphafold2 gerbil prestin structure (Q9JKQ2 · S26A5_MERUN), which are closely related to that of the mouse. The unresolved gap in the structure at residues 159-164 was filled with the corresponding residues in the Alphafold structure after the two models had been superimposed at residues 138-158 and 165-194. Residues 581-613 in the STAS domain, which are not resolved in any structures, predicted to be unstructured, and not reliably modeled by Alphafold, were omitted from our simulation system. The resulting model of wildtype (WT) gerbil prestin was set up in a 1-Palmitoyl-2-oleoyl-*sn*-glycero-3-phosphocholine (POPC) and cholesterol bilayer with a 9:1 ratio and surrounding solvent with a NaCl concentration of 150 mM using the CHARMM-GUI V1.7 web interface (Jo et al., 2008; Lee et al., 2016). In a similar fashion, two models of the S396E and S398E mutant were created by using the “mutate residue” functionality in CHARMM-GUI. All models were then prepared for MD simulation in the same manner as described next. We employed the CHARMM27 force field with cmap for proteins (MacKerell et al., 2004), CHARMM36 for lipids (Klauda et al., 2010) and the CHARMM TIP3P water model. All titratable residues were simulated in their default protonation states at pH7, as predicted by PROPKA 3.1 (Olsson et al., 2011). Relaxation and equilibration were performed with GROMACS (Abraham et al., 2015; Páll et al., 2020) following the CHARMM-GUI protocol (Jo et al., 2008) with an initial energy minimization stage and six stages of equilibration during which harmonic restraints on protein and lipids were successively reduced to zero. Simulations were performed at 296 K.

#### All-atom, explicit solvent Anton MD simulations

Production runs of WT and the S398E mutant were performed on the Anton 2 supercomputer, a special-purpose computer for molecular MD simulations of biomolecules (Shaw et al., 2014). The system’s charge was neutralized with the addition of sodium and chloride ions. Additional sodium and chloride ions were added to produce a free NaCl concentration of 150 mM in solution. The CHARMM36 (MacKerell et al., 1998; MacKerell et al., 2004; Best et al., 2012) force field was used for proteins and ions, and the TIP3P model was used for waters. The RESPA algorithm (Tuckerman et al., 1992) was used to integrate the long-range nonbonded forces every 7.5 fs and short-range nonbonded and bonded forces every 2.5 fs. Long-range electrostatic forces were calculated using the k-Gaussian split Ewald method (Shan et al., 2005). The lengths of bonds to hydrogen atoms were held fixed using the nSHAKE algorithm. The simulations were run at 296K and 1 atm using the Nosé-Hoover chain thermostat (Martyna et al., 1992) and the Martyna-Tobias-Klein (Martyna et al., 1994) barostat, respectively. The RESPA algorithm and temperature and pressure controls were implemented using the multigrator scheme (Lippert et al., 2013) with a time step of 2.5 fs. Temperatures were maintained using Nosé-Hoover chains with time constants τ_*T*_ = 0.042 s. Semi-isotropic pressure coupling using Nosé-Hoover chains kept the system at 1 atm across both temperature groups with time constants τ_*P*_ = 0.042 s. Coordinates were written to trajectory files every 240 ps.

Simulations were either carried out under equilibrium conditions (0 mV membrane potential) or set up to model a strong negatively polarized membrane of about –150 mV by using the constant electric field approximation (Roux, 2008). A static external electric field in the z direction (a 3-vector electric field [0, 0, -0.024] in unit kcal/(mol·Å·e) which is equivalent to -0.00104 V/Å) was applied to the system. Under the constant electric field approximation, the potential drop across the membrane equals the average length of the simulation cell in field direction (145 Å) times the field strength, resulting in approximately –150 mV. In order to avoid sudden changes to the system, the electric field was linearly increased from zero to its maximum value over the initial 200 ns of the simulation; after 200 ns the electric field remained constant for the rest of the simulation.

Anton2 enables the simulation of relatively long times, so here we report 5 µs of trajectory data for WT (at 0 mV and –150 mV) and 2.5 µs for S398E (at 0 mV and –150 mV).

#### All-atom, explicit solvent GROMACS MD simulations

Production runs of prestin with the S396E mutation were performed on Arizona State University’s *sol* supercomputer (Jennewein et al., 2023) with GROMACS version 2024.2 (Abraham et al., 2015; Páll et al., 2020). The system’s charge was neutralized with the addition of sodium and chloride ions. Additional sodium and chloride ions were added to produce a free NaCl concentration of 150 mM in solution. As for the Anton2 simulations, the CHARMM36 (MacKerell et al., 1998; MacKerell et al., 2004; Best et al., 2012) force field was used for proteins and ions, and the CHARMM TIP3P model was used for waters. The stochastic velocity rescaling thermostat (Bussi and Parrinello, 2007) was used with a time constant of 1ps and three separate temperature-coupling groups for protein, lipids, and solvent. The Parrinello–Rahman barostat (Parrinello and Rahman, 1981) with a time constant of 5 ps and a compressibility 4.5 × 10^−5^bar^-1^ was used for semi-isotropic pressure coupling. Coulomb interactions were calculated with the fast-smooth particle-mesh Ewald (SPME) method (Essmann et al., 1995) with an initial real-space cutoff of 1.2 nm, which was optimized by the Gromacs GPU code at run time, and interactions beyond the cutoff were calculated in reciprocal space with a fast Fourier transform on a grid with 0.12 nm spacing and fourth-order spline interpolation. The Lennard– Jones forces were switched smoothly to zero between 1.0 and 1.2 nm and the potential was shifted over the whole range and decreased to zero at the cutoff. Bonds to hydrogen atoms were converted to rigid holonomic constraints with the P-LINCS algorithm (Hess, 2008). The classical equations of motions were integrated with the leapfrog algorithm with a time step of 2 fs.

To reproduce the same equilibrium and negatively polarized conditions as in the Anton2 setup, simulations were either carried out at 0 mv membrane potential or with a constant electric field E = – 0.00103 V/Å applied to the system, resulting in a membrane potential of approximately –150 mV. Three repeats for each membrane potential were simulated, totaling more than 1.3 µs each (0 mV: 492.3 ns, 414.9 ns, 466.2 ns; –150 mV: 479.9 ns, 483.1 ns, 459.9 ns).

#### Analysis

Trajectory analysis was carried out with Python scripts based on MDAnalysis (Michaud-Agrawal et al., 2011; Gowers et al., 2016). The radial distribution function (RDF) of Cl^-^ was calculated for heavy atoms (C, N, O) using the MDAnalysis tool *interRDF*. The RDFs showed that the first shell of ligand atoms of Cl^-^ in the binding site always lies within a 5 Å radius. Therefore, binding of Cl^-^ ions to the prestin binding site was assessed with the following distance criterion: any Cl^-^ ion within 5 Å of any heavy atom (carbon, nitrogen, oxygen) of residue 137, 396, 397, 398 was considered bound. Molecular images were prepared UCSF Chimera (Pettersen et al., 2004).

## Results

**Figure 1 A and B** show the frequency responses of control mouse cNLC in membrane patches. The response is like that of guinea pig prestin in that real and imaginary components of cNLC display differing frequency response, but similar bell-shaped voltage dependence. The absolute magnitude of cNLC,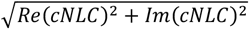, near V_h_ (**Fig. 1B**), displays a stretched-exponential roll-off typical of many physical materials, including viscoelastic behavior of glasses (Langer, 2012), cytoskeleton (Lieleg et al., 2011) and others that must overcome a multitude of energy barriers. For prestin, it indicates that sensor charge movement works against viscoelastic loads within the membrane whose influence is frequency dependent. The average cut-off frequency (F_is_) is 27.34 +/-4.31 kHz for the mouse, indicated by the intersection of real and imaginary components, which in a simple 2-state voltage-only Boltzmann model identifies the -3 dB point of absolute cNLC (see Appendix in (Santos-Sacchi and Tan, 2022)). Being stretched-exponential in form, not 2-state Lorentzian, the actual corresponding reduction in Abs(cNLC) at F_is_ is near 90% of half-magnitude. We previously found guinea pig patch F_is_ to be about 19 kHz (Santos-Sacchi et al., 2023). We cannot appropriately compare mouse and GP kinetics because OHCs were obtained from different best frequency regions of the animals’ cochleas.

We previously showed that chloride anions and salicylate can alter prestin kinetics (Santos-Sacchi and Song, 2016; Santos-Sacchi and Tan, 2019). Chloride binding to the identified binding pocket in prestin is not required for the generation of NLC since negatively charged mutations of coordinating residues in the pocket (e.g., residues 396, 398 and 399) can substitute for chloride (Gorbunov et al., 2014; Gorbunov et al., 2018; Butan et al., 2022). Anions have no influence on NLC in that case, presumably because they are not bound. **Fig. 2** shows the results of MD simulations of two such gerbil prestin mutation, S396E and S398E, which abrogate anion binding (Gorbunov et al., 2014; Gorbunov et al., 2018; Oliver et al., 2019; Butan et al., 2022). In the normal binding pocket, the chloride anion readily interacts with the coordinating residues, whereas with either mutation binding is absent or systematically reduced. What effect does a mutation that removes chloride binding have on hearing and cNLC frequency response? We use a KI mouse where the mutation S396E has been engineered into mouse prestin to evaluate this question.

**Figure 2.**
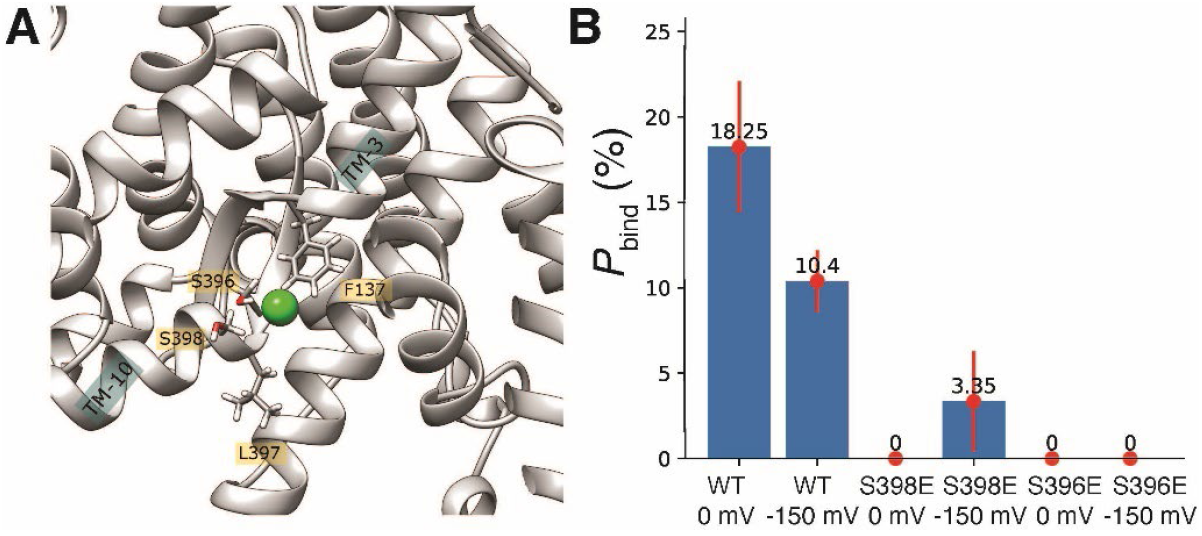
MD simulations show absence of binding in gerbil prestin with mutation S396E and S398E. **A)** Snapshot of the anion binding site of WT prestin with bound Cl^−^ ion (green) between helices TM-3 and TM-10 and residues F132, S396, L397, S398 (highlighted). **B)** Binding probability for Cl^−^ (fraction of time that a Cl^−^ ion occupied the binding site during the simulation, measured in percent) for WT, S398E and S396E at 0 mV and –150 mV, reported as an average over the independent binding probabilities observed for protomer A and B. The full error bar represents the absolute value of the difference between the A and B value and is an approximate measure of the variability of obtaining the binding probability from the MD simulations; for WT and S398E, the binding probabilities for A and B were obtained from long multi-microsecond simulations, for S396E, binding probabilities were averaged over three shorter repeat simulations, all of which showed 0% binding.

**Fig. 3** shows that hearing in this KI mouse is normal, both ABR and distortion product emissions being no different from control mice. As might be expected from this result, **Fig. 4** shows that in OHCs high frequency charge movements in control and S396E mice are the same (27.34 +/-4.31 kHz vs. 29.72 +/-1.67 kHz; t-test, 2-tail: p=0.625). The maintained stretched-exponential behavior in S396E mouse OHCs indicates that the viscoelastic membrane environment dominates the behavior of prestin kinetics at high frequencies. Under whole-cell conditions, altering intracellular chloride results in shifts in V_h_ (Rybalchenko and Santos-Sacchi, 2003a; Santos-Sacchi et al., 2006b; Rybalchenko and Santos-Sacchi, 2008), increases causing a negative shift. Because the S396E mutation effectively saturates the chloride binding site, compared to our controls we might expect a V_h_ shift to negative values. Indeed, a negative shift in V_h_ relative to controls did occur (−99.5 +/-7.5 mV vs. -127.8 +/-7.0 mV; t-test, 2-tail: p=0.033); however, see Discussion.

**Figure 3.**
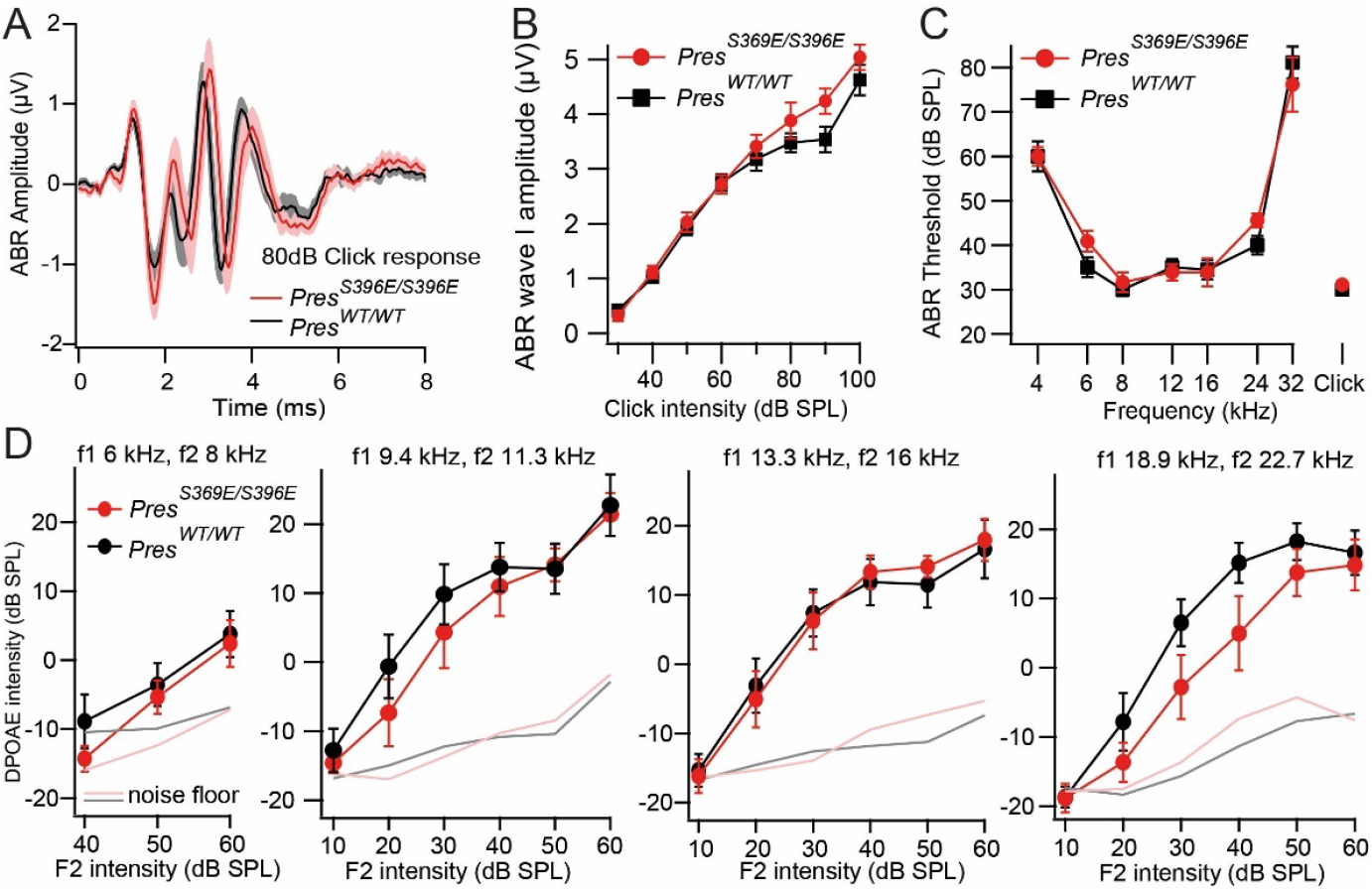
Normal hearing function in *Pres*^*S396E/S396E*^ mutant mice. **A-C**, *Pres*^*S396E/S396E*^ mutants (red, n=9) had normal ABR waveforms (**A**), wave I amplitude growth functions (measured from peak to trough, **B**) and thresholds compared wildtype littermates (black, n=9) at the age of 13 weeks. Some thresholds at 32 kHz exceeded the maximal loudspeaker output of 80 dB SPL and were set to 90 dB SPL for calculation of the average. **D)** DPOAE amplitude growth functions for different frequency combinations were comparable different *Pres*^*S396E/S396E*^ mutants (red, n=7) and wildtype littermates (black, n=7) at the age of 4 weeks. All data are displayed as means ±SEM.

**Figure 4.**
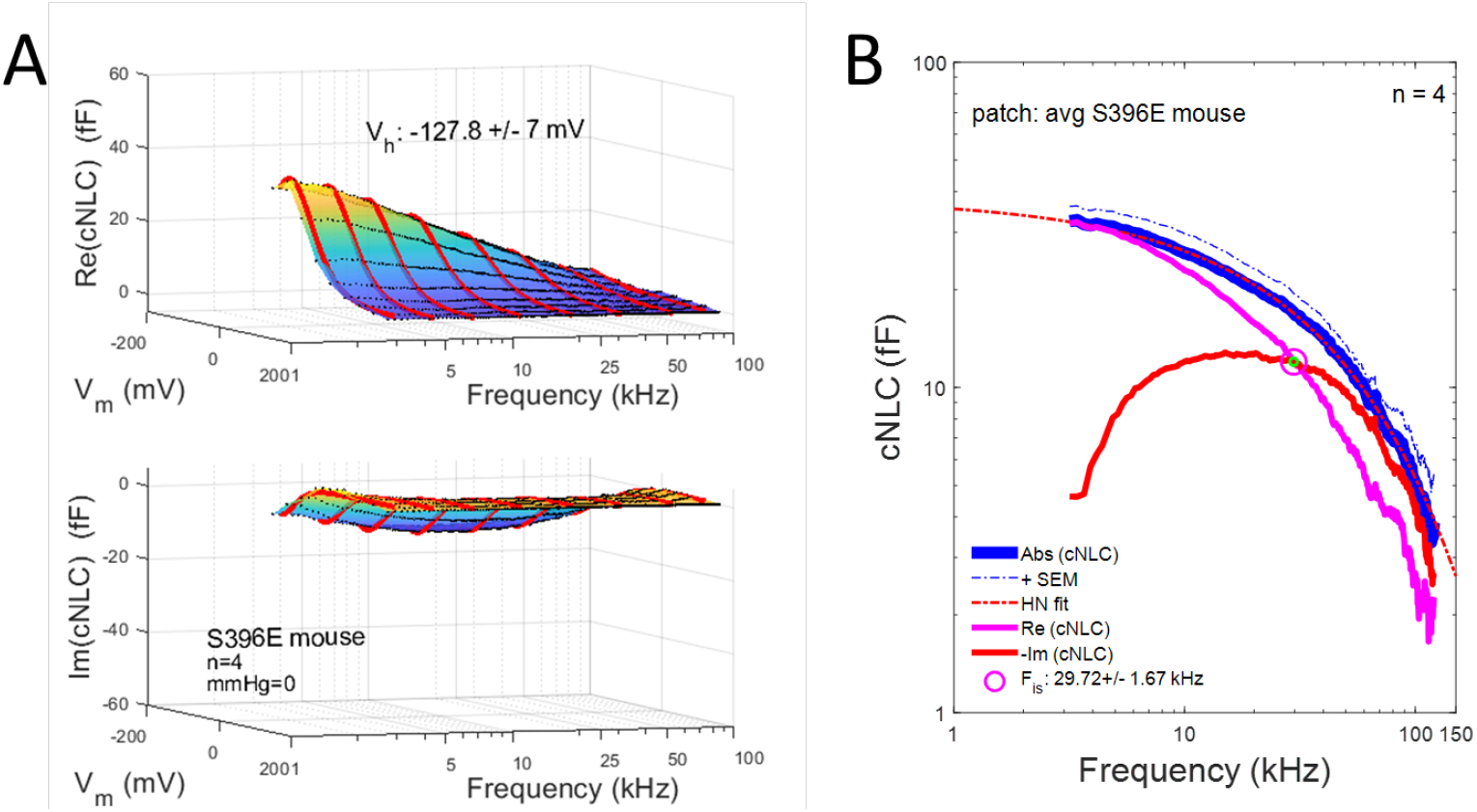
Average complex NLC in S396E mouse OHC membrane patches. **A)** Real (top panel) and imaginary (bottom panel) components of cNLC across frequency and membrane voltage. Black lines indicate data and red lines are Boltzmann fits at selected frequencies. Averages (SE) @ 5.12 kHz V_h_ : -127.8 (7.0) mV; Q_max_: 5.4 (0.7) fC; *z*: 0.58 (0.01). V_h_ is significantly left-shifted compared to b6 mouse control OHCs. Note that real component decreases with frequency while imaginary component increases then decreases with frequency. No pipette pressure is delivered. **B)** Average frequency response of cNLC components near V_h_. Abs (cNLC) decreases across frequency in a stretched-exponential manner, indicated by the HN pink dotted line fit. The contribution of the imaginary component to absolute magnitude dominates at the highest frequencies. Average F_is_ (magenta circle) determined from individual recordings is 29.72 (1.67) kHz, not different from control b6 mouse OHCs. Green point marker indicates intersection of averaged traces at 29.7 kHz.

Finally, we compared the tension dependence of S396E and WT NLC in membrane patches, positive tension causing a depolarizing shift in WT V_h_. (Iwasa, 1993; Kakehata and Santos-Sacchi, 1995). We previously showed that anion binding affinity decreases when tension is applied to the membrane (Song and Santos-Sacchi, 2010), and we suggested that piezoelectric-like behavior in prestin, namely, V_h_ shifts due to tension, results from allosteric modulation of prestin by anions. In the absence of anion binding in the S396E mutant, we reasoned that piezoelectric-like behavior would be absent. This is not the case, however, as – 8 mmHg pressure in the patch pipette shifts V_h_ of WT and S396E mutant equally; shifts are 14.9 +/-2.5 mV vs. 11.3 +/-0.6 mV, respectively; n=3, t-test, 2-tail: p=0.24. Thus, other molecular mechanisms must be responsible for the mechanical sensitivity of prestin.

## Discussion

Prestin works mechanically into the ultrasonic range to enhance our ability to perceive sound through a process termed “cochlear amplification”. The kinetics of prestin underlie its ability to influence cochlear mechanics at these high frequencies (Santos-Sacchi and Tan, 2018; Santos-Sacchi, 2019). Beyond intramolecular constraints within prestin itself, the frequency responsiveness of prestin depends on the impinging loads within the membrane bilayer, as well as other cellular/environmental loads (Santos-Sacchi et al., 2019). We previously showed that possible loads associated with the macro-patch technique and membrane tension have only slight effects on NLC frequency response (Santos-Sacchi and Tan, 2022), so our results here reflect characteristic performance of prestin in the mouse induced by experimental perturbations directly to prestin.

We find that prestin frequency response in normal mice surpasses that we previously measured in the guinea pig (Santos-Sacchi et al., 2023), although we are unable to unambiguously compare mouse and guinea pig kinetics because OHCs were obtained from different best frequency regions of each animal. Given that caveat, mouse NLC cut-off frequency, characterized by F_is_, where real and imaginary components of cNLC intersect, is about 8 kHz greater than in guinea pig (27 vs. 19 kHz). Considering that the frequency roll-off in prestin in each species is stretched exponential in nature rather than Lorentzian, F_is_ corresponds to a frequency where there is greater than half magnitude reduction in total charge movement, both in and out of phase with voltage, quantified by Abs(cNLC). Consequently, based on our fitting for the control b6 mouse (**Fig. 1 B**), Abs(cNLC) is down at 80 kHz by 14.73 dB. For the guinea pig we previously found greater roll-off, Abs(cNLC) being down at 80 kHz by 17.71 (Santos-Sacchi et al., 2023).

### Chloride binding and NLC

Chloride anions can affect prestin kinetics (Santos-Sacchi and Song, 2016; Santos-Sacchi and Tan, 2022), with mutations altering binding site anion access being influential (Takahashi et al., 2016; Takahashi et al., 2023). Although it had been speculated that Cl^-^ acts as an extrinsic voltage sensor (Oliver et al., 2001), instead anions likely foster allosteric-like modulation of prestin activity and kinetics (Rybalchenko and Santos-Sacchi, 2003b; Rybalchenko and Santos-Sacchi, 2003a, 2008; Song and Santos-Sacchi, 2010, 2013; Santos-Sacchi and Song, 2016). Binding pocket mutations are confirmatory (e.g., residues 396, 398, and 399), since they enable voltage dependency of prestin without anion presence (Gorbunov et al., 2014; Gorbunov et al., 2018; Oliver et al., 2019; Butan et al., 2022). Our MD simulations confirm that the S396E and S398E mutations of gerbil prestin halt chloride entry and residence within its anion binding pocket (**Fig. 2**). Though we had suggested that the allosteric action of anions on prestin could underlie its piezoelectric-like behavior (Song and Santos-Sacchi, 2010), we show here that even in the absence of chloride binding modulation by tension as occurs in WT prestin, the S396E mutant is as tension-sensitive as WT prestin.

Our frequency response measures of S396E NLC indicate no influence of the mutation, F_is_ being no different than that of controls (see above). Based on our fits for the S396E mouse, Abs(cNLC) is down at 80 kHz by 14.56 dB. This is essentially the same as the control results. However, we find that the S396E mutation appears to alter V_h_ in a predictable manner. As we mentioned above, under whole-cell conditions, reducing intracellular chloride results in reversible positive shifts in V_h_ (Rybalchenko and Santos-Sacchi, 2003a; Santos-Sacchi et al., 2006b; Rybalchenko and Santos-Sacchi, 2008). Given that the S396E mutation essentially mimics saturation of the chloride binding site, a significant V_h_ shift to negative values arises compared to our controls. This negative shift was also found in S396E prestin transfected cells [see supplement in (Takahashi et al., 2023)]; however, in that paper another similar mutation (S396D) did not shift V_h_. Further investigation on the V_h_ shift in anion binding pocket mutations is warranted.

Interestingly, Takahashi et al. (Takahashi et al., 2023), chose to use the anion-insensitive mutation S396D for their KI mouse model, which like our S396E KI model has normal hearing. Their reasoning was based on using a mutation (S396D) whose V_h_ was close to WT prestin, rather than shifted markedly in the hyperpolarized direction (S396E). It had been thought that large operating point shifts away from WT prestin V_h_ will disrupt cochlear amplification, this concept being the rationale for the “499” KI mouse (Dallos et al., 2008), where V_h_ was shifted far to positive potentials. The fact that we find normal hearing with a mutation that has a large negative shift in voltage operation point indicates that the shift itself was not the reason for impaired hearing in the “499” mouse. Indeed, subsequently the “499” mouse was shown to have impaired fast kinetics, which likely results in poor hearing (Homma et al., 2013). It is thus significant that our S396E KI mouse OHCs exhibit normal motor kinetics, despite a large V_h_ shift, thereby preserving normal hearing. Accordingly, a fast frequency response of prestin charge movement appears to be a key determinant for normal hearing.

Finally, we note that prestin tertiary structures of all species studied to-date are essentially indistinguishable (Bavi et al., 2021; Ge et al., 2021; Butan et al., 2022; Futamata et al., 2022), yet it is obvious that mammals have evolved to utilize a wide range of species-specific auditory frequency bandwidths. Recently, Shi and colleagues (Liu et al., 2014; Liu et al., 2018; Liu et al., 2022) have entertained the concept that prestin primary structure in echolocating mammals may have evolved to alter its voltage sensitivity, their rationale being based partially on our measures of the NLC Boltzmann *z* parameter (an indicator of voltage sensitivity) across the frequency spectrum in guinea pig OHCs (Santos-Sacchi et al., 1998). In fact, such evolutionary pressures apparently have occurred to enhance high frequency hearing in echolocating species (Liu et al., 2014; Liu et al., 2018; Liu et al., 2022). Paradoxically, however, the slope of prestin charge vs. voltage is much shallower in species of augmented high frequency hearing, indicating that the maximum gain of OHC **eM** at V_h_ (δ eM_Vh_/δ Vm_Vh_) in these species is less than in non-echolocating species. Why such a reduction in OHC mechanical activity would occur is perplexing. Structural studies and frequency measures of these prestin constructs may be informative. In any case, we hypothesize that, in the absence of chloride influence on high frequency performance, any special auditory requirements of individual species for cochlear amplification may have additionally evolved to reduce impinging loads on the protein, including those within the membrane (Santos-Sacchi and Tan, 2022) and organ of Corti.

## Acknowledgements

This research was supported by NIH-NIDCD R56DC021057, R01 DC016318 and R01 DC008130. Anton 2 computer time was provided by the Pittsburgh Supercomputing Center (PSC) under allocation MCB180088P through Grant R01GM116961 from the National Institutes of Health; the Anton 2 machine at PSC was generously made available by D.E. Shaw Research. Additionally supported by the German Research Foundation through Collaborative Research Center 889 project A06 to N.S.

## References

Abraham MJ, Murtola T, Schulz R, Páll S, Smith JC, Hess B, Lindahl E (2015) GROMACS: High performance molecular simulations through multi-level parallelism from laptops to supercomputers. SoftwareX 1:19–25.

Alverez F, Alegria A, Colmenero J (1991) Relationship between the time-domain Kohlrausch-Williams-Watts and frequency-domain Havriliak-Negami relaxation functions. Physical Review B 44:7306–7312.

Bavi N, Clark MD, Contreras GF, Shen R, Reddy BG, Milewski W, Perozo E (2021) The conformational cycle of prestin underlies outer-hair cell electromotility. Nature 600:553–558.

Best RB, Zhu X, Shim J, Lopes PEM, Mittal J, Feig M, Mackerell J, Alexander D (2012) Optimization of the additive CHARMM all-atom protein force field targeting improved sampling of the backbone $\phi$, $\psi$ and side-chain $\chi_1$ and $\chi_2$ dihedral angles. J Chem Theory Comput 8:3257–3273.

Brownell WE, Bader CR, Bertrand D, de Ribaupierre Y (1985) Evoked mechanical responses of isolated cochlear outer hair cells. Science 227:194–196.

Bussi G, Parrinello M (2007) Accurate sampling using Langevin dynamics. Physical Review E—Statistical, Nonlinear, and Soft Matter Physics 75:056707.

Butan C, Song Q, Bai J-P, Tan WJT, Navaratnam D, Santos-Sacchi J (2022) Single particle cryo-EM structure of the outer hair cell motor protein prestin. Nat Commun 13:290.

Dallos P, Santos-Sacchi J, Flock A (1982) Intracellular recordings from cochlear outer hair cells. Science 218:582–584.

Dallos P, Wu X, Cheatham MA, Gao J, Zheng J, Anderson CT, Jia S, Wang X, Cheng WH, Sengupta S, He DZ, Zuo J (2008) Prestin-based outer hair cell motility is necessary for mammalian cochlear amplification. Neuron 58:333–339.

Dong X, Ehrenstein D, Iwasa KH (2000) Fluctuation of motor charge in the lateral membrane of the cochlear outer hair cell. Biophys J 79:1876–1882.

Essmann U, Perera L, Berkowitz ML, Darden T, Lee H, Pedersen LG (1995) A smooth particle mesh Ewald method. The Journal of chemical physics 103:8577–8593.

Fernandez JM, Bezanilla F, Taylor RE (1982) Distribution and kinetics of membrane dielectric polarization. II. Frequency domain studies of gating currents. JGenPhysiol 79:41–67.

Futamata H, Fukuda M, Umeda R, Yamashita K, Tomita A, Takahashi S, Shikakura T, Hayashi S, Kusakizako T, Nishizawa T, Homma K, Nureki O (2022) Cryo-EM structures of thermostabilized prestin provide mechanistic insights underlying outer hair cell electromotility. Nat Commun 13:6208.

Gale JE, Ashmore JF (1994) Charge displacement induced by rapid stretch in the basolateral membrane of the guinea-pig outer hair cell. Proc R Soc Lond B Biol Sci 255:243–249.

Gale JE, Ashmore JF (1997) An intrinsic frequency limit to the cochlear amplifier. Nature 389:63–66.

Ge J, Elferich J, Dehghani-Ghahnaviyeh S, Zhao Z, Meadows M, von Gersdorff H, Tajkhorshid E, Gouaux E (2021) Molecular mechanism of prestin electromotive signal amplification. Cell 184:4669–4679 e4613.

Geertsma ER, Oliver D (2023) SLC26 Anion Transporters. Handb Exp Pharmacol.

Gorbunov D, Hartmann J, Renigunta V, Oliver D (2018) A glutamate scan identifies an electrostatic switch for prestin activity. Midwinter Meeting Abstracts of the Associatio for Research in Otolaryngology.

Gorbunov D, Sturlese M, Nies F, Kluge M, Bellanda M, Battistutta R, Oliver D (2014) Molecular architecture and the structural basis for anion interaction in prestin and SLC26 transporters. Nat Commun 5:3622.

Gowers RJ, Linke M, Barnoud J, Reddy TJE, Melo MN, Seyler SL, Dotson DL, Domański J, Buchoux S, Kenney IM, Beckstein O (2016) MDAnalysis: A Python package for the Rapid Analysis of Molecular Dynamics Simulations. In: Proceedings of the 15th Python in Science Conference (Benthall S, Rostrup S, eds), pp 98--105. Austin, TX.

Hess B (2008) P-LINCS: A parallel linear constraint solver for molecular simulation. J Chem Theory Comput 4:116–122.

Homma K, Duan C, Zheng J, Cheatham MA, Dallos P (2013) The V499G/Y501H mutation impairs fast motor kinetics of prestin and has significance for defining functional independence of individual prestin subunits. J Biol Chem 288:2452–2463.

Humphrey W, Dalke A, Schulten K (1996) VMD -- Visual Molecular Dynamics. J Molecular Modelling 14:33--38.

Iwasa KH (1993) Effect of stress on the membrane capacitance of the auditory outer hair cell. Biophys J 65:492–498.

Jennewein DM, Lee J, Kurtz C, Dizon W, Shaeffer I, Chapman A, Chiquete A, Burks J, Carlson A, Mason N (2023) The Sol Supercomputer at Arizona State University. In: Practice and Experience in Advanced Research Computing, pp 296–301.

Jing Z, Rutherford MA, Takago H, Frank T, Fejtova A, Khimich D, Moser T, Strenzke N (2013) Disruption of the presynaptic cytomatrix protein bassoon degrades ribbon anchorage, multiquantal release, and sound encoding at the hair cell afferent synapse. J Neurosci 33:4456–4467.

Jo S, Kim T, Iyer VG, Im W (2008) CHARMM-GUI: a web-based graphical user interface for CHARMM. J Comput Chem 29:1859–1865.

Kakehata S, Santos-Sacchi J (1995) Membrane tension directly shifts voltage dependence of outer hair cell motility and associated gating charge. Biophys J 68:2190–2197.

Klauda JB, Venable RM, Freites JA, O’Connor JW, Tobias DJ, Mondragon-Ramirez C, Vorobyov I, MacKerell J, Alexander D, Pastor RW (2010) Update of the CHARMM all-atom additive force field for lipids: validation on six lipid types. J Phys Chem B 114:7830–7843.

Langer JS (2012) Shear-transformation-zone theory of viscosity, diffusion, and stretched exponential relaxation in amorphous solids. Phys Rev E Stat Nonlin Soft Matter Phys 85:051507.

Lee J, Cheng X, Swails JM, Yeom MS, Eastman PK, Lemkul JA, Wei S, Buckner J, Jeong JC, Qi Y, Jo S, Pande VS, Case DA, Brooks r, Charles L, MacKerell J, Alexander D, Klauda JB, Im W (2016) CHARMM-GUI Input Generator for NAMD, GROMACS, AMBER, OpenMM, and CHARMM/OpenMM Simulations Using the CHARMM36 Additive Force Field. J Chem Theory Comput 12:405–413.

Lieleg O, Kayser J, Brambilla G, Cipelletti L, Bausch AR (2011) Slow dynamics and internal stress relaxation in bundled cytoskeletal networks. Nat Mater 10:236–242.

Lippert RA, Predescu C, Ierardi DJ, Mackenzie KM, Eastwood MP, Dror RO, Shaw DE (2013) Accurate and efficient integration for molecular dynamics simulations at constant temperature and pressure. J Chem Phys 139:164106.

Liu Z, Qi FY, Zhou X, Ren HQ, Shi P (2014) Parallel sites implicate functional convergence of the hearing gene prestin among echolocating mammals. Mol Biol Evol 31:2415–2424.

Liu Z, Qi FY, Xu DM, Zhou X, Shi P (2018) Genomic and functional evidence reveals molecular insights into the origin of echolocation in whales. Sci Adv 4:eaat8821.

Liu Z, Chen P, Xu DM, Qi FY, Guo YT, Liu Q, Bai J, Zhou X, Shi P (2022) Molecular convergence and transgenic evidence suggest a single origin of laryngeal echolocation in bats. iScience 25:104114.

MacKerell A et al. (1998) All-atom empirical potential for molecular modeling and dynamics studies of proteins. JPhysChemB 102:3586--3616.

MacKerell J A. D.,, Feig M, Brooks III CL (2004) Extending the treatment of backbone energetics in protein force fields: limitations of gas-phase quantum mechanics in reproducing protein conformational distributions in molecular dynamics simulations. JCompChem 25:1400--1415.

Martyna GJ, Klein ML, Tuckerman M (1992) Nosé–Hoover chains: The canonical ensemble via continuous dynamics. The Journal of Chemical Physics 97:2635–2643.

Martyna GJ, Tobias DJ, Klein ML (1994) Constant pressure molecular dynamics algorithms. J Chem Phys 101:4177–4189.

Mason MJ, Simpson AK, Mahaut-Smith MP, Robinson HP (2005) The interpretation of current-clamp recordings in the cell-attached patch-clamp configuration. Biophys J 88:739–750.

Michaud-Agrawal N, Denning EJ, Woolf TB, Beckstein O (2011) MDAnalysis: A Toolkit for the Analysis of Molecular Dynamics Simulations. J Comp Chem 32:2319--2327.

Oliver D, Gorbunov D, Hartmann J, Lenz D, Renigunta V (2019) An electrostatic switch for gating the electromechanical activity of SLC26A5 (prestin). Biophys J 116:169a.

Oliver D, He DZ, Klocker N, Ludwig J, Schulte U, Waldegger S, Ruppersberg JP, Dallos P, Fakler B (2001) Intracellular anions as the voltage sensor of prestin, the outer hair cell motor protein. Science 292:2340–2343.

Olsson MHM, Søndergaard CR, Rostkowski M, Jensen JH (2011) PROPKA3: Consistent Treatment of Internal and Surface Residues in Empirical $\mathrmpK_\mathrma$ Predictions. J Chem Theory Comput 7:525–537.

Páll S, Zhmurov A, Bauer P, Abraham M, Lundborg M, Gray A, Hess B, Lindahl E (2020) Heterogeneous parallelization and acceleration of molecular dynamics simulations in GROMACS. The Journal of Chemical Physics 153.

Parrinello M, Rahman A (1981) Polymorphic transitions in single crystals: A new molecular dynamics method. Journal of Applied physics 52:7182–7190.

Pettersen EF, Goddard TD, Huang CC, Couch GS, Greenblatt DM, Meng EC, Ferrin TE (2004) UCSF Chimera--a visualization system for exploratory research and analysis. J Comput Chem 25:1605–1612.

Roux B (2008) The membrane potential and its representation by a constant electric field in computer simulations. BiophysJ 95:4205--4216.

Russell IJ, Sellick PM (1978) Intracellular studies of hair cells in the mammalian cochlea. JPhysiol (Lond) 284:261–290.

Rybalchenko V, Santos-Sacchi J (2003a) Cl-flux through a non-selective, stretch-sensitive conductance influences the outer hair cell motor of the guinea-pig. J Physiol 547:873–891.

Rybalchenko V, Santos-Sacchi J (2003b) Allosteric modulation of the outer hair cell motor protein prestin by chloride. In: Biophysics of the Cochlea: From Molecules to Models (Gummer A, ed), pp 116–126. Singapore: World Scientific Publishing.

Rybalchenko V, Santos-Sacchi J (2008) Anion control of voltage sensing by the motor protein prestin in outer hair cells. Biophys J 95:4439–4447.

Santos-Sacchi J (1991) Reversible inhibition of voltage-dependent outer hair cell motility and capacitance. J Neurosci 11:3096–3110.

Santos-Sacchi J (2019) The speed limit of outer hair cell electromechanical activity. HNO 67:159–164.

Santos-Sacchi J, Dilger JP (1988) Whole cell currents and mechanical responses of isolated outer hair cells. Hear Res 35:143–150.

Santos-Sacchi J, Navarrete E (2002) Voltage-dependent changes in specific membrane capacitance caused by prestin, the outer hair cell lateral membrane motor. Pflugers Arch 444:99–106.

Santos-Sacchi J, Song L (2016) Chloride anions regulate kinetics but not voltage-sensor Qmax of the solute carrier SLC26a5. Biophys J 110:1–11.

Santos-Sacchi J, Tan W (2018) The Frequency Response of Outer Hair Cell Voltage-Dependent Motility Is Limited by Kinetics of Prestin. J Neurosci 38:5495–5506.

Santos-Sacchi J, Tan W (2019) Voltage Does Not Drive Prestin (SLC26a5) Electro-Mechanical Activity at High Frequencies Where Cochlear Amplification Is Best. iScience 22:392–399.

Santos-Sacchi J, Tan W (2022) On the frequency response of prestin charge movement in membrane patches. Biophys J May 20;S0006-3495(22)00413-1. .

Santos-Sacchi J, Iwasa KH, Tan W (2019) Outer hair cell electromotility is low-pass filtered relative to the molecular conformational changes that produce nonlinear capacitance. J Gen Physiol 151:1369–1385.

Santos-Sacchi J, Navaratnam D, Tan WJT (2021) State dependent effects on the frequency response of prestin’s real and imaginary components of nonlinear capacitance. Sci Rep 11:16149.

Santos-Sacchi J, Bai JP, Navaratnam D (2023) Megahertz Sampling of Prestin (SLC26a5) Voltage-Sensor Charge Movements in Outer Hair Cell Membranes Reveals Ultrasonic Activity that May Support Electromotility and Cochlear Amplification. J Neurosci 43:2460–2468.

Santos-Sacchi J, Song L, Zheng J, Nuttall AL (2006a) Control of mammalian cochlear amplification by chloride anions. J Neurosci 26:3992–3998.

Santos-Sacchi J, Song L, Zheng JF, Nuttall AL (2006b) Control of mammalian cochlear amplification by chloride anions. J Neurosci 26:3992–3998.

Santos-Sacchi J, Kakehata S, Kikuchi T, Katori Y, Takasaka T (1998) Density of motility-related charge in the outer hair cell of the guinea pig is inversely related to best frequency. Neurosci Lett 256:155–158.

Shan Y, Klepeis JL, Eastwood MP, Dror RO, Shaw DE (2005) Gaussian split Ewald: A fast Ewald mesh method for molecular simulation. J Chem Phys 122:54101.

Shaw DE et al. (2014) Anton 2: Raising the Bar for Performance and Programmability in a Special–Purpose Molecular Dynamics Supercomputer. In: SC ‘14: Proceedings of the International Conference for High Performance Computing, Networking, Storage and Analysis, pp 41–53.

Song L, Santos-Sacchi J (2010) Conformational state-dependent anion binding in prestin: evidence for allosteric modulation. Biophys J 98:371–376.

Song L, Santos-Sacchi J (2013) Disparities in voltage-sensor charge and electromotility imply slow chloride-driven state transitions in the solute carrier SLC26a5. Proc Natl Acad Sci U S A.

Takahashi S, Santos-Sacchi J (2001) Non-uniform mapping of stress-induced, motility-related charge movement in the outer hair cell plasma membrane. Pflugers Arch 441:506–513.

Takahashi S, Cheatham MA, Zheng J, Homma K (2016) The R130S mutation significantly affects the function of prestin, the outer hair cell motor protein. J Mol Med (Berl) 94:1053–1062.

Takahashi S, Zhou Y, Kojima T, Cheatham MA, Homma K (2023) Prestin’s fast motor kinetics is essential for mammalian cochlear amplification. Proc Natl Acad Sci U S A 120:e2217891120.

Tuckerman M, Berne BJ, Martyna GJ (1992) Reversible multiple time scale molecular dynamics. The Journal of Chemical Physics 97:1990–2001.

Zheng J, Shen W, He DZ, Long KB, Madison LD, Dallos P (2000) Prestin is the motor protein of cochlear outer hair cells. Nature 405:149–155.

